# Imp is required for timely exit from quiescence in *Drosophila* type II neuroblasts

**DOI:** 10.1101/2022.07.15.500194

**Authors:** Jordan A. Munroe, Mubarak H. Syed, Chris Q. Doe

## Abstract

Stem cells must balance proliferation and quiescence, with excess proliferation favoring tumor formation, and premature quiescence preventing proper organogenesis. *Drosophila* brain neuroblasts are a model for investigating neural stem cell entry and exit from quiescence. Neuroblasts begin proliferating during embryogenesis, enter quiescence prior to larval hatching, and resume proliferation 12-30h after larval hatching. Here we focus on the mechanism used to exit quiescence, focusing on the “type II” neuroblasts. There are 16 type II neuroblasts in the brain, and they undergo the same cycle of embryonic proliferation, quiescence, and proliferation as do most other brain neuroblasts. We focus on type II neuroblasts due to their similar lineage as outer radial glia in primates (both have extended lineages with intermediate neural progenitors) and because of the availability of specific markers for type II neuroblasts and their progeny. Here we characterize the role of Insulin-like growth factor II mRNA-binding protein (Imp) in type II neuroblast proliferation and quiescence. Imp has previously been shown to promote proliferation in type II neuroblasts, in part by acting antagonistically to another RNA-binding protein called Syncrip (Syp). Here we show that reducing Imp levels delays neuroblast exit from quiescence in type II neuroblasts, acting independently of Syp, with Syp levels remaining low in both quiescent and newly proliferating type II neuroblasts. We conclude that Imp promotes exit from quiescence, a function closely related to its known role in promoting neuroblast proliferation.

## Introduction

The generation of neuronal diversity is essential for proper brain assembly and function. This is particularly true for the primate cortex, which derives from a specialized neural stem cell called outer radial glia (oRG). These stem cells are thought to have driven cortical expansion and diversity during evolution (1–3), but how they regulate their proliferation remains incompletely understood.

One way to help understand oRG lineages is to use model organisms that contain neural stem cells with lineages similar to oRGs, which can be used to generate testable hypotheses for investigating primate oRG lineages. In *Drosophila*, there is a small pool of 16 neural stem cells in the brain (eight stem cells per brain lobe), called type II neuroblasts (TIINBs), that undergo a lineage similar to primate oRGs to generate neurons (4–6) (Figure 1A). In primates these oRGs generate neurons of the cortex; in *Drosophila* the TIINBs generate neurons of the adult central complex (CX), a region important for navigation, sleep, and sensorimotor integration (7). Like oRGs, TIINBs undergo repeated asymmetric divisions to produce a series of transit amplifying cells called Intermediate Neural Progenitor (INPs), which themselves undergo a more limited division pattern to generate a series of ganglion mother cells (GMCs) which undergo a single terminal division to produce pairs of neurons and/or glia (Figure 1A, left)(4–6).

**Figure 1.**
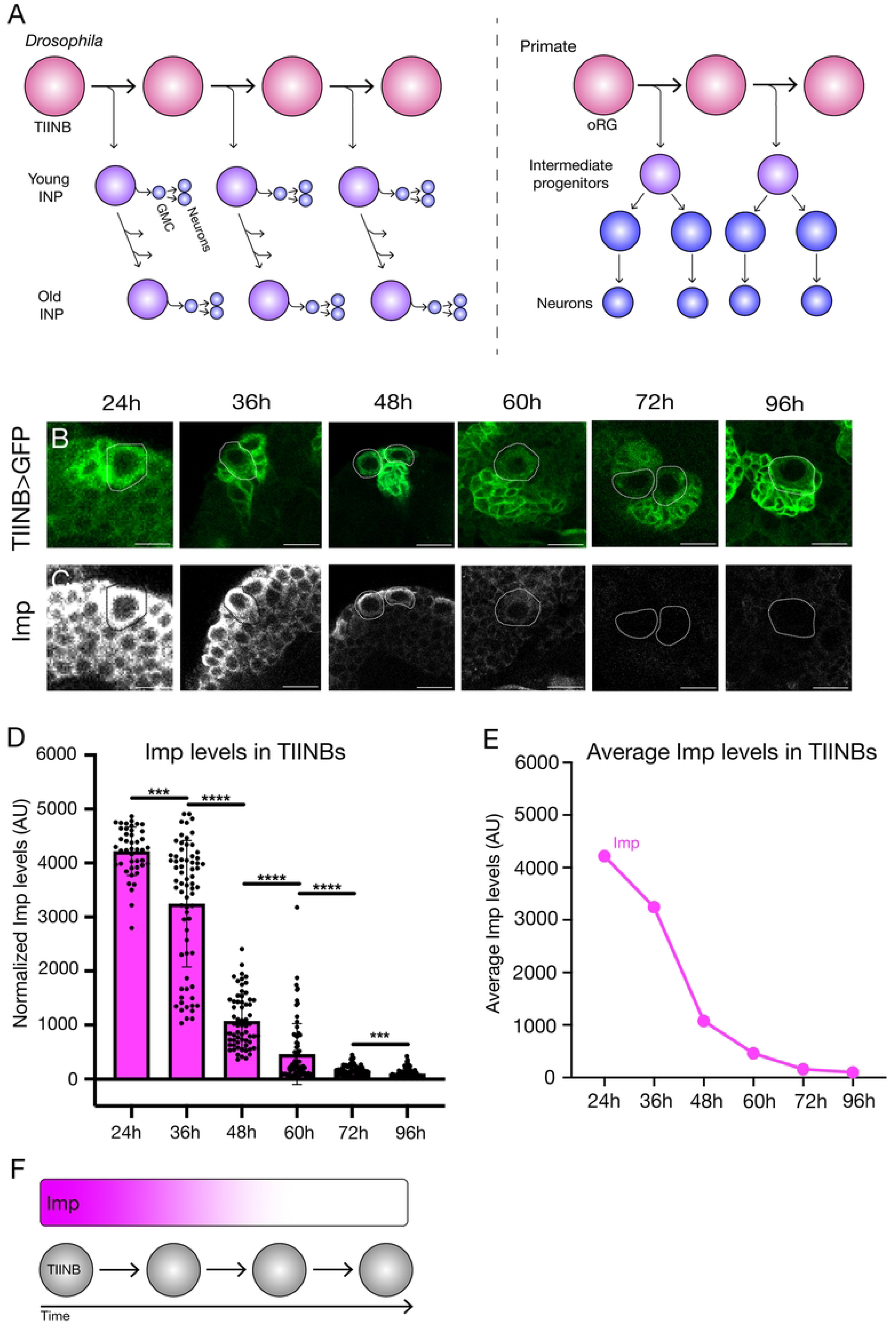
Quantification of the Imp gradient in type II neuroblasts. (A) Type II neuroblast lineage (left) (4–6) and outer radial glial lineage (right), adapted from (27). (B,C) Imp protein forms a high-to-low gradient in type II neuroblasts during larval life (hours are time after larval hatching in this and following figures). Type II neuroblasts are identified by expression of *pointed-gal4 UAS-GFP.* Scale bar, 20 μm. (D,E) Quantification of Imp protein levels (see methods for details) for all n’s (D) or for the average levels (E). (F) Summary.

Neuronal diversity is generated at each step in the TIINB lineage. TIINBs change gene expression over time as they generate distinct INPs, with some genes limited to early lineage expression such as insulin-like growth factor II mRNA-binding protein (Imp), Chinmo, and Lin-28; other genes are only expressed late in the lineage such as Syncrip (Syp), Broad, and E93 (8,9). These genes are called candidate temporal transcription factors (TTFs) or temporal identity factors due to their potential role in specifying different neuronal fates based on their time of birth. Subsequently, each individual INP undergoes a TTF cascade to generate molecularly distinct GMCs (10–12). Thus, the TIINBs appear to be an excellent model for understanding oRG lineages in primates.

Another important aspect of TIINB lineages is how their pattern of proliferation is regulated to generate large populations of neurons without tumorigenesis. TIINBs begin their lineage in the embryonic brain, followed by a period of quiescence at the transition from embryo to first larval instar (L1), and then proliferation resumes between 12-30 hours after larval hatching (13,14); subsequently all times refer to hours after larval hatching. This is similar to the pattern of proliferation-quiescence-proliferation exhibited by most other embryonic larval neuroblast lineages (15,16). Previous work has shown that neuroblast quiescence is achieved through the accumulation of nuclear Prospero (Pros) (16,17), and upon exit from quiescence each TIINB will generate ~60 INPs that produce hundreds of neurons and glia throughout larval development (4–6,18–20). Previous work has shown that low Notch signaling activity is required to drive mushroom body neuroblasts (MB NBs) out of quiescence, while Syp recruits the mediator complex and Pros to drive the MB NBs into decommissioning (21,22). This terminal exit from the cell cycle is also driven by the loss of proliferation and differentiation due to low Imp expression (21,22). High Imp expression in early larval life promotes neuroblast proliferation via the stabilization of *myc* and *chinmo* RNAs as well as inhibition of the Mediator complex (9,21,23). This makes Imp an attractive candidate for studying how TIINBs initiate exit from quiescence. Here we focus on the role of Imp in regulating neuroblast proliferation in TIINB lineages, where we identify a novel role for Imp in promoting TIINB exit from quiescence.

## Results

### Type II neuroblasts exhibit a high-to-low Imp protein gradient overtime

Previous work has shown that Imp forms a high-to-low RNA and protein gradient in all assayed neuroblast populations (23), but at just a few timepoints. Here we used *Pointed-gal4 (pnt-gal4)* crossed to *UAS-GFP* to identify TIINBs and co-stained for Imp at 12h intervals throughout larval stages, from 24h to 96h after larval hatching; note that all times subsequently refer to hours after larval hatching (Figure 1B). We found that Imp protein forms a gradient from high to low over the first 60h of larval life, becoming virtually undetectable from 72-96h (Figure 1B-F). We conclude that Imp levels drop continuously in TIINBs during larval life.

### ImpRNAi and Imp overexpression have opposing effects on the timing of the Imp protein gradient in type II neuroblasts

To alter the Imp protein gradient, we performed Imp RNAi in TIINBs. We used *pnt-gal4 UAS-impRNAi* to reduce Imp protein levels specifically in TIINB lineages. We found that ImpRNAi in TIINBs significantly reduced Imp protein levels, although an Imp protein gradient persisted, effectively shifting the Imp gradient to earlier times in development (Figure 2A,C-E). In contrast, overexpression of Imp within TIINB lineages results in higher levels of Imp, without abolishing its gradient, effectively shifting the Imp gradient to later times in development (Figure 2B-E). We conclude that ImpRNAi or Imp overexpression reduces or increases Imp protein levels, respectively, and thus they are effective tools for manipulating Imp protein levels in TIINBs.

**Figure 2.**
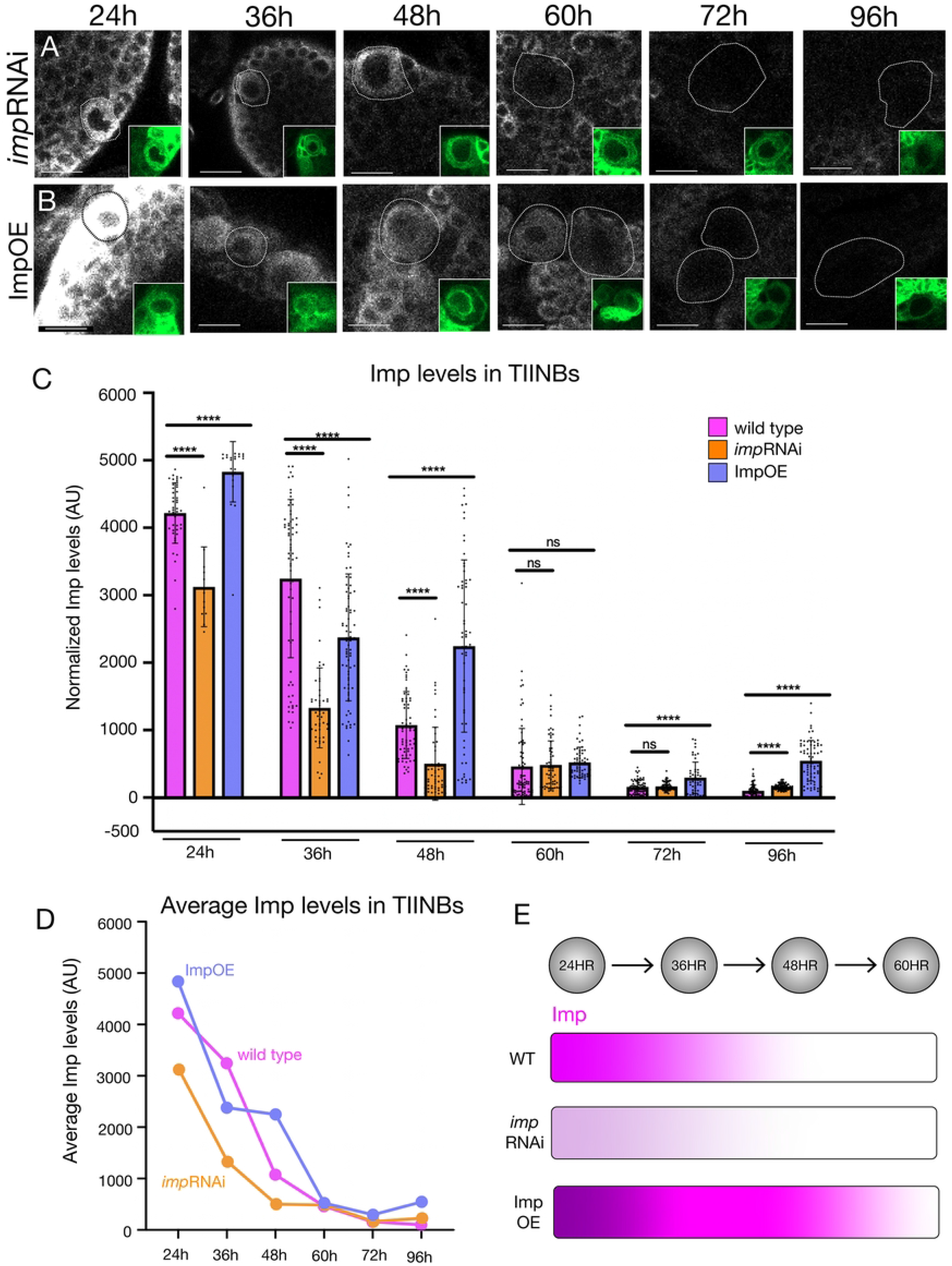
Imp RNAi and Imp overexpression result in reduced or increased Imp protein levels. Wild type Imp levels are shown in Figure 1. (A) Imp RNAi within type II neuroblasts (inset: *pointed-gal4 UAS-GFP*) leads to lower Imp levels without disrupting the protein gradient; quantified in C. Scale bar, 20 μm. (C) Imp overexpression within type II neuroblasts (inset: *pointed-gal4 UAS-GFP*) leads to higher Imp levels without disrupting the protein gradient; quantified in C. Scale bar, 20 μm. (C, D) Quantification of Imp protein levels in type II neuroblasts in wild type, Imp RNAi, and Imp overexpression. (C) Histogram showing all n’s; (D) graph showing average values. (E) Summary.

### *pnt-gal4 UAS-GFP* can be used to selectively label proliferating type II neuroblasts

Imp has been shown to promote neuroblast proliferation, and the decline in Imp levels in late larva contributes to termination of neuroblast proliferation (21,22). Here we asked a related question: does reduction in Imp levels in TIINB delay exit from quiescence? Proliferating versus quiescent TIINBs can be distinguished by Deadpan (Dpn) and Cyclin E (CycE) expression: proliferative neuroblasts are Dpn+CycE+ whereas quiescent neuroblasts are Dpn+CycE- (15,16). We found that *pnt-Gal4 UAS-GFP* was only expressed by proliferating TIINBs (Figure 3A; quantified in 3C), and no quiescent neuroblasts expressed *pnt-Gal4 UAS-GFP* (Figure 3B; quantified in 3C). This allowed us to quantify how many of the 16 TIINBs were proliferating, and infer the remainder were quiescent (see below). We conclude that *pnt-gal4 UAS-GFP* can be used to identify proliferating TIINBs (Figure 3D).

**Figure 3.**
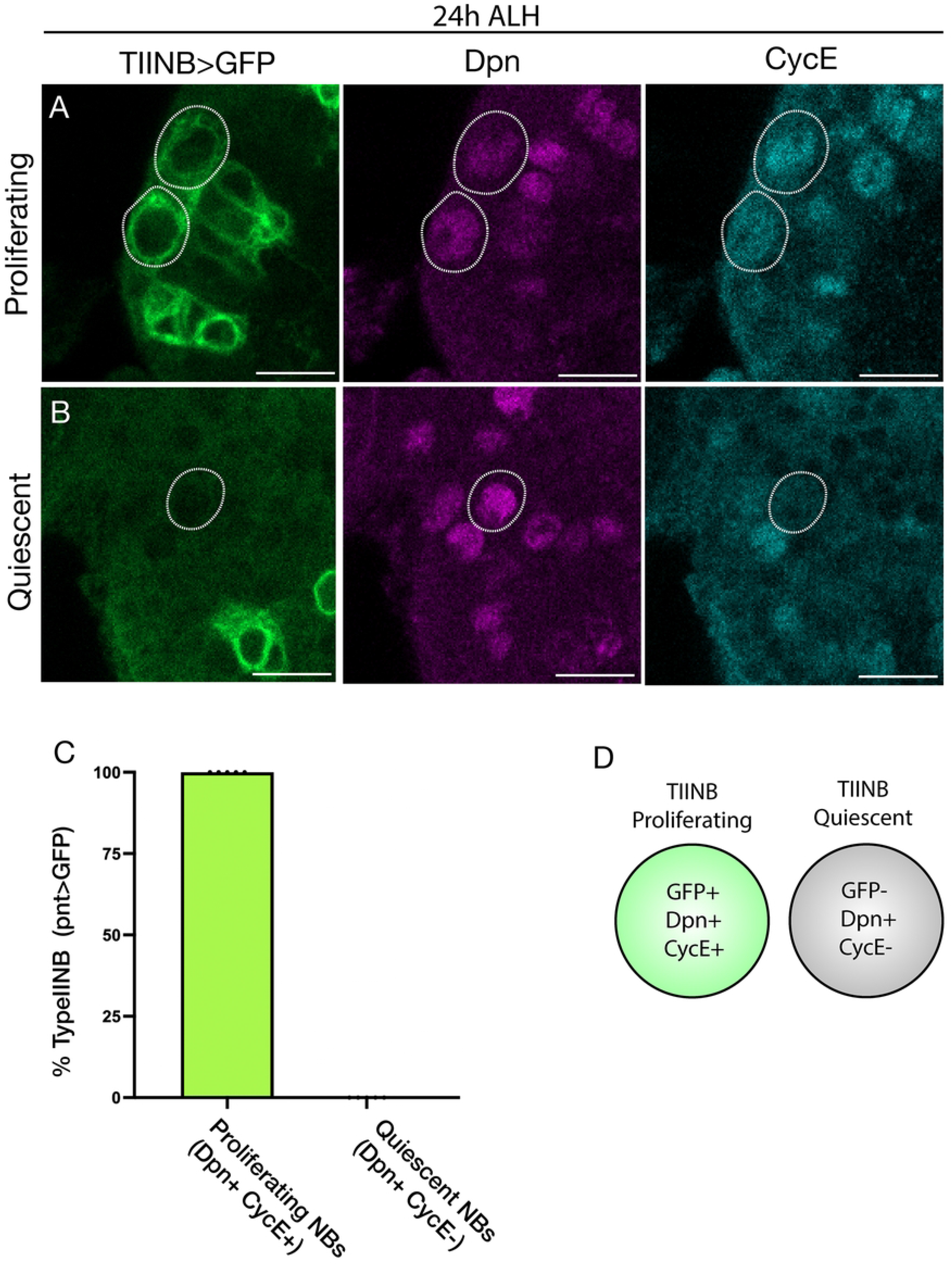
*Pointed-gal4 UAS-GFP+* TIINBs have exited quiescence and are proliferative. (A) Type II neuroblasts are circled and identified by *pointed-gal4 UAS-GFP* (green), Dpn (magenta), and reconfirmed as proliferative by CycE (cyan) at 24h. Scale bar is 5 μm. (B) *pointed-gal4 UAS-GFP* (green) and CycE (cyan) are not expressed in quiescent type II neuroblasts, but Dpn (magenta) is still present. Quiescent cells are circled. Scale bar is 5 μm. (C) Histogram of cells that are Dpn+. One hundred percent of type II neuroblasts that are positive for GFP *(pointed-gal4 UAS-GFP)* are Dpn+ and CycE+, while 0% of cells that are GFP-, Dpn+, CycE-. (D) Summary.

### Imp is required for timely exit from quiescence in type II neuroblasts

High Imp expression early in larval development promotes neuroblast proliferation, while late, low Imp expression leads to neuroblast decommissioning (21,22). We wanted to know if high Imp expression early in larval life promoted TIINB exit from quiescence. To answer this question, we decreased Imp levels specifically in TIINB lineages and quantified the number of proliferating TIINBs at intervals from 24h to 96h. We used *pnt-gal4* to *UAS-GFP* to identify proliferating TIINBs, *UAS-ImpRNAi* (to reduce Imp levels), and Dpn to mark all neuroblasts (proliferating or quiescent). In wild type, at 24h ~8 of the 16 TIINBs are *pnt-Gal4 UAS-GFP+* and thus have exited quiescence, with the remainder still in quiescence. By 36h, all 16 TIINBs have exited quiescence and are proliferative (Figure 4A,B). In contrast, following ImpRNAi, only ~2 TIINBs have exited quiescence at 24h, and it takes until 72h for all 16 TIINBs to exit quiescence and become proliferative (Figure 4A,B). We also wanted to see if ImpRNAi delayed exit from quiescence in specific TIINB lineages – e.g. the pair of lateral DL neuroblasts or dorsomedial DM neuroblasts – but each class had an indistinguishable time of exit from quiescence (data not shown). We conclude that Imp promotes exit from quiescence in TIINBs.

**Figure 4.**
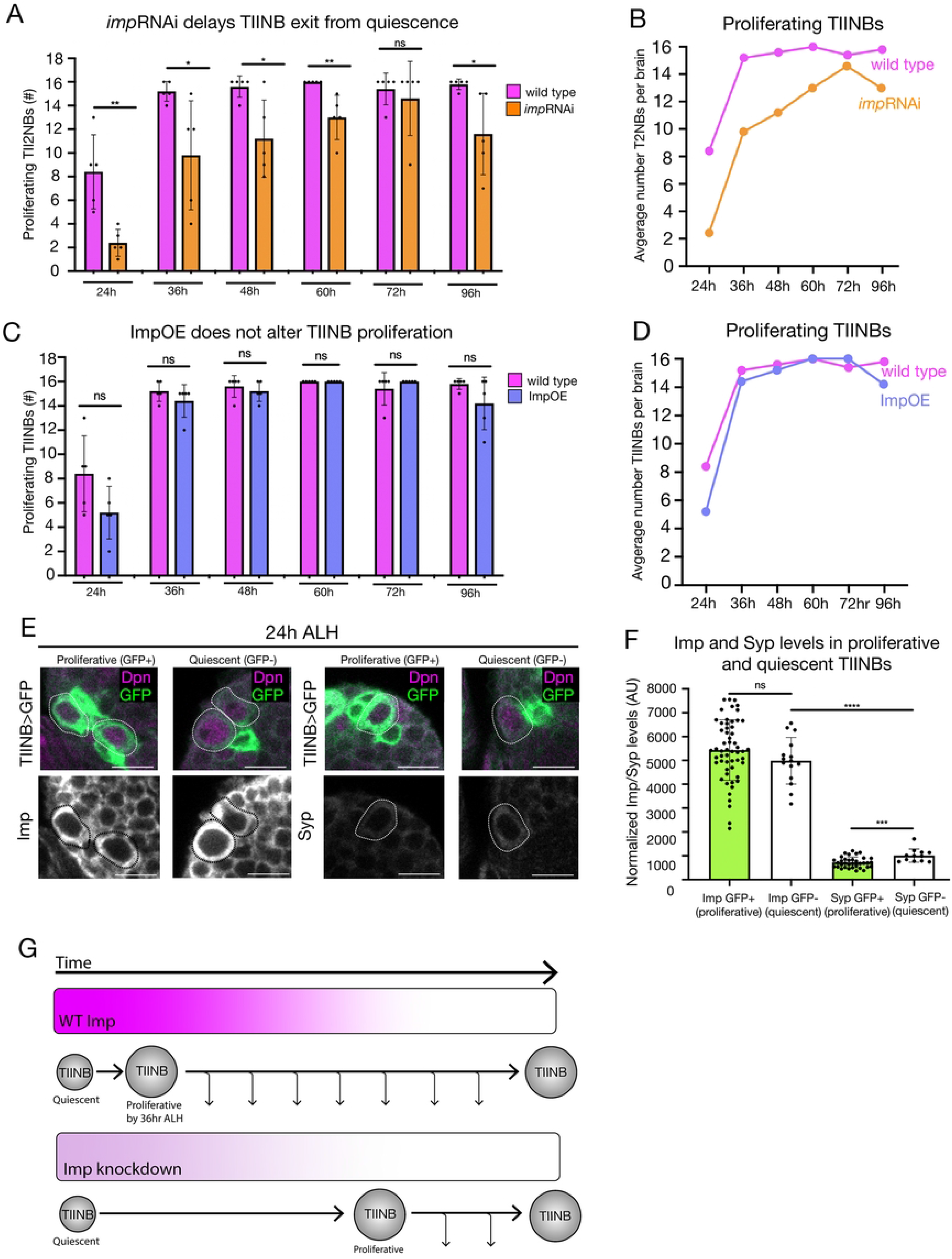
Imp is required for timely exit from quiescence in type II neuroblasts. (A,B) Quantification of proliferating type II neuroblast numbers (expressing *Pointed-gal4 UAS-GFP)* over larval life in wild type and Imp RNAi. Note that there is a maximum of 16 type II neuroblasts per brain. In wild type, all neuroblasts have exited quiescence/resumed proliferating by 36h as shown by *Pointed-gal4 UAS-GFP* expression. Imp RNAi delays exit from quiescence and the full complement of 16 proliferating type II neuroblasts is not achieved until 72h as shown by Pointed-gal4 UAS-GFP expression. (C,D) Quantification of proliferating type II neuroblast numbers (*Pointed-gal4 UAS-GFP+*) across larval development for wild type and Imp overexpression. There is no difference in exit from quiescence between wild type and Imp overexpression genotypes. (E) Imp and Syp levels are the same in quiescent and proliferating type II neuroblasts. Proliferating type II neuroblasts (circled; first and third columns) are identified by expression of *pointed-gal4 UAS-GFP* (green), Dpn (magenta), and lack of Asense (not shown). Quiescent type II neuroblasts do not express *pointed-gal4 UAS-GFP* (green) but can be identified as Dpn+ (magenta) and lack of Asense (data not shown). (F) Quantification of Imp and Syp levels in quiescent and proliferating type II neuroblasts at 24h. (G) Summary.

To determine if higher levels of Imp could drive precocious exit from quiescence, we used *pointed-gal4* to drive *UAS-Imp* specifically in TIINB lineages. This manipulation results in significantly more Imp protein in TIINBs (Figure 2), but overexpression of Imp does not induce precocious exit from quiescence in TIINBs (Figure 4C,D). We conclude that Imp is necessary but not sufficient to drive TIINB exit from quiescence.

Because Imp promotes exit from quiescence, we asked whether quiescent TIINBs have low Imp and proliferating TIINBs have high Imp levels. Interestingly, we observed comparable levels of Imp in proliferating TIINBs (Figure 4E, first column; quantified in 4F) and quiescent TIINBs (Figure 4E, second column; quantified in 4F). Because Imp and Syp can cross-repress each other (23), we assayed Syp levels in proliferating and quiescent TIINBs. As expected, we found Syp to be expressed at a lower level than Imp in both proliferating and quiescent TIINBs (Figure 4E, third and fourth columns; quantified in 4F). Interestingly, Syp levels in quiescent TIINBs were slightly higher than Syp levels in proliferative TIINBs (Figure 4F), showing a correlation between higher Syp levels and neuroblast quiescence. We conclude that Imp is expressed in quiescent neuroblasts and is necessary but not sufficient for timely exit from quiescence (Figure 4G).

## Discussion

It is well documented in previous studies that Imp is expressed in a temporal gradient in many central brain neuroblasts (8,9,22,23). In this study we have confirmed the Imp gradient in TIINBs from 24h – 96h and have quantified Imp levels in wild type as well as after ImpRNAi knockdown or Imp overexpression. While both knockdown and overexpression show significant changes in Imp levels, the Imp gradient is maintained throughout larval life in all cases. Interestingly, at 36h Imp overexpression levels are lower than WT control levels, but only at this timepoint. This suggests a post-transcriptional ‘homeostatic’ mechanism that reduces Imp levels when they are experimentally increased. A possible explanation for this is Imp targeting by microRNA *let-7. let-7* targets Imp in *Drosophila* male testis (24) and is present in MB NBs where it targets the temporal transcription factor Chinmo, which is also present in TIINBs (25). Thus, *let-7* may regulate Imp in TIINBs and should be explored in future work.

At 24h wild type larval brains show ~8-10 TIINBs active, and all 16 TIINBs (8 neuroblasts per brain lobe) are active and proliferating by 36h. Imp knockdown results in only ~2-4 TIINBs at 24h and all 16 TIINBs are not proliferating until 72h. This late exit from quiescence shows that Imp is necessary for timely exit from quiescence. Previous studies have shown that high levels of Imp in TIINBs are required to maintain large neuroblast size and proliferative activity through the stabilization of *myc* RNA (22), and overexpression of Imp in neuroblasts can extend proliferation (21,22). Our results add to these findings by showing that Imp is required for TIINB timely exit from quiescence. Additionally, Imp knock down in TIINBs promotes early exit from cell cycle at the end of larval life (21). Imp overexpression in TIINBs did not change the rate at which TIINBs exit from quiescence. Thus, Imp is necessary but not sufficient for exit from quiescence. These findings suggest that a minimum level of Imp is required for the exit from quiescence. A potential mechanism for this would be a negative feedback loop driven by over-expression of Imp, which could lead to over-proliferation if not regulated. Again, a candidate factor for regulation of Imp levels as TIINBs exit quiescence is *let-7.*

We quantified Imp levels in both quiescent and proliferative TIINBs to see how they varied and saw no change. We also wanted to compare Syp levels to Imp levels in quiescent and proliferative TIINBs. Syp is required for the entrance into quiescence and decommissioning (21), but it was unknown what Syp levels are in TIINBs nearing the end of quiescence early in larval life. We compared Syp levels in proliferating TIINBs to quiescent TIINBs but found that Syp levels were significantly lower than Imp levels, consistent with their cross-repressive regulation. Interestingly, Syp levels in quiescent TIINBs were higher than Syp levels in proliferative TIINBs, showing a correlation between high Syp levels and neuroblast quiescence, and consistent with earlier work showing Syp is required to elevate levels of nuclear Prospero and enter quiescence (21).

## Materials and Methods

### Key Resource Table

| Reagent | Designation | Source | Identifiers | Additional information |
| --- | --- | --- | --- | --- |
| Species<br>(*D melanogaster*) | Pointed-Gal4 | (26) | n/a | TIINB driver |
| Species<br>(*D melanogaster*) | UAS-ImpRNAi | BDSC | #34977 | Imp knockdown |
| Species<br>(*D melanogaster*) | UAS-Imp | Syed Lab (UNM) | Macdonald lab | Imp overexpression |
| Species<br>(*D melanogaster*) | UAS-myr::GFP | BDSC | #32198 | Membrane bound GFP under UAS control |
| Antibody, polyclonal | Chicken anti-GFP | Abcam (Eugene, OR) | n/a | 1:1000 |
| Antibody, polyclonal | Rabbit anti-Imp | MacDonald lab (UT Austin) | n/a | 1:1000 |
| Antibody, polyclonal | Rabbit anti-Syp | Desplan Lab (NYU) | n/a | 1:1000 |
| Antibody, polyclonal | Rat anti-Dpn | Abcam (Eugene, OR) | n/a | 1:20 |
| Antibody, polyclonal | Rabbit anti-CycE | Santa Cruz Biotechnology | #C1209 | 1:500 |
| Antibody, polyclonal | Guinea pig anti-Asense | Wang lab (Duke) | n/a | 1:500 |
| Antibody, polyclonal | Secondary antibodies | Thermofisher (Eugene, OR) | n/a | 1:400, 1:200 (Dpn only) |

| Fly genotype used in each figure | Figure | Synopsis |
| --- | --- | --- |
| ; UAS-myr::GFP ; Pnt-Gal4 | 1, 2, 3 | Control. GFP = TIINBs |
| ; UAS-myr::GFP ; Pnt-Gal4 X ; ; UAS ImpRNAi | 2, 3 | Imp RNAi. GFP = TIINBs |
| UAS-Imp ; Sco/Cyo ; X ; UAS-myr::GFP ; Pnt Gal4 | 2, 3 | Imp overexpression. GFP = TIINBs. |

#### Antibodies and immunostaining

All larvae were raised at 25°C and dissected in Hemolymph Like buffer 3.1 (HL3.1) (NaCl 70mM, KCl 5mM, CaCl_2_ 1.5mM, MgCl_2_ 4mM, sucrose 115mM, HEPES 5mM, NaHCO_3_ 10mM, and Trehalose 5mM in double distilled water). Larvae were grown to specified time points, dissected, mounted on poly-D-lysine coated slips (Neuvitro, Camas, WA), and incubated for 30 minutes in 4% paraformaldehyde solution in Phosphate Buffered Saline (PBS) with 1% Triton-X (1% PBS-T) at room temperature. Larval brains were washed twice with 0.5% PBS-T and incubated for 1-7 days at 4°C in a blocking solution of 1% goat serum (Jackson ImmunoResearch, West Grove, PA), 1% donkey serum (Jackson ImmunoResearch, West Grove, PA), 2% dimethyl sulfoxide in organosulfur (DMSO), and 0.003% bovine serum albumin (BSA) (Fisher BioReagents, Fair Lawn, NJ Lot #196941). Larval brains were incubated overnight at 4°C in a solution of primary antibodies (see Key Resource Table) in 0.5% PBS-T. Larval brains were washed for at least 60 minutes in 0.5% PBS-T at room temperature, and then incubated overnight at 4°C in a solution of secondary antibodies (see Key Resource Table) in 0.5% PBS-T. Brains were washed in 0.5% PBS-T for at least 60 minutes at room temperature. Brains were dehydrated by going through a series of 10-minute washes in 30%, 50%, 70%, and 90% EtOH, and two rounds of 10 minutes in 100% EtOH and two rounds of 10 minutes in xylene (MP Biomedicals, LLC, Saolon, OH, Lot# S0170), then mounted in dibutyl phthalate in xylene (DPX; Sigma-Aldrich, cat. no. 06522). Brains sat in DPX for at least 48 hours at 4°C or 72 hours (48 hours at room temperature and 24 hours at 4°C) before imaging.

#### Imaging and statistical analysis

All Imp data were collected with identical confocal settings; all Syp data were collected with identical confocal settings. Fluorescent images were collected on Zeiss LSM 800. TIINBs were counted using the cell counter plugin in FIJI (https:://imagej.net/software/fiji/). Imp pixel density in each TIINB was calculated in FIJI. TIINBs were manually selected in FIJI using the polygon lasso tool, and the area and Raw Integrated Density (RID) was measured. The nucleus of each TIINB went through the same analysis steps. Imp levels were normalized to cell area using the equation: (Cell Body^RID^ – Nucleus^RID^) / (Cell Body^Area^ – Nucleus^Area^). Two-tailed student t-tests were used to compare two sets of data. *p<0.05; **p<0.01; ***p<0.001; ****p<0.0001. All graphs and statistical analysis were done in Prism (GraphPad Software, San Diego, CA).

#### Figure production

Images for figures were taken in FIJI. Figures were assembled in Adobe Illustrator (Adobe, San Jose, CA). Any changes in brightness or contrast were applied to the entire image.

## Acknowledgements

We thank Noah Dillon and Gonzalo Morales Chaya for comments on the manuscript, and Adam Fries for help with developing a fluorescent analysis method.

## Ethical Approval and Consent to participate

n/a

## Consent for publication

All authors consent for this to be published

## Availability of data and material

No new data sets or fly genotypes were developed.

## Authors’ contributions

JM performed all the experiments, generated all the figures, and wrote the first draft of the paper. MHS and CQD supervised the project and edited figures and text.

## Competing Interests

None.

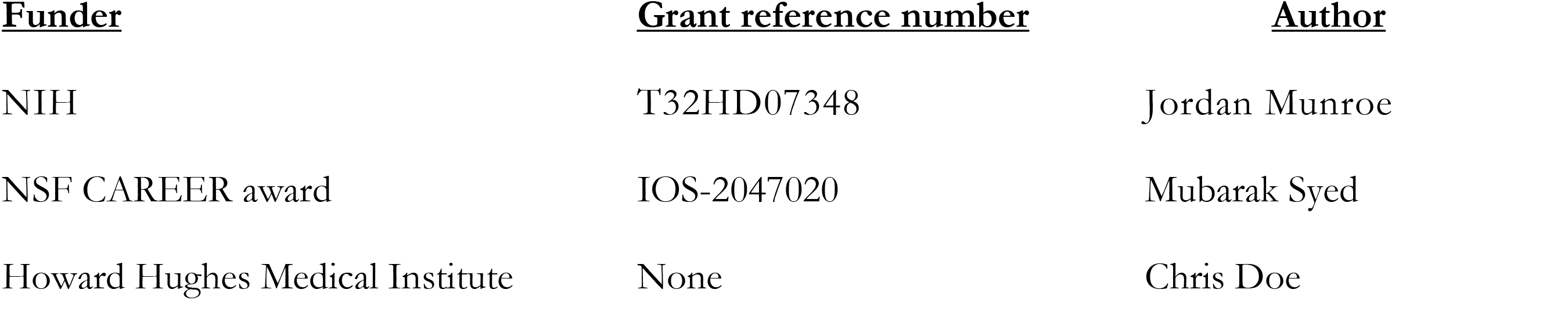

| <u>Funder</u> | <u>Grant reference number</u> | <u>Author</u> |
| --- | --- | --- |
| NIH | T32HD07348 | Jordan Munroe |
| NSF CAREER award | IOS-2047020 | Mubarak Syed |
| Howard Hughes Medical Institute | None | Chris Doe |

The funders had no role in study design, data collection and interpretation, or the decision to submit the work for publication.

## References

1. Hansen DV, Lui JH, Parker PR, Kriegstein AR. Neurogenic radial glia in the outer subventricular zone of human neocortex. Nature. 2010 Mar 25;464(7288):554–61.

2. Fietz SA, Kelava I, Vogt J, Wilsch-Brauninger M, Stenzel D, Fish JL, et al. OSVZ progenitors of human and ferret neocortex are epithelial-like and expand by integrin signaling. Nat Neurosci. 2010 Jun;13(6):690–9.

3. Noctor SC, Martinez-Cerdeno V, Ivic L, Kriegstein AR. Cortical neurons arise in symmetric and asymmetric division zones and migrate through specific phases. Nat Neurosci. 2004 Feb;7(2):136–44.

4. Boone JQ, Doe CQ. Identification of Drosophila type II neuroblast lineages containing transit amplifying ganglion mother cells. Dev Neurobiol. 2008 Aug;68(9):1185–95.

5. Bowman SK, Rolland V, Betschinger J, Kinsey KA, Emery G, Knoblich JA. The tumor suppressors Brat and Numb regulate transit-amplifying neuroblast lineages in Drosophila. Dev Cell. 2008 Apr;14(4):535–46.

6. Bello BC, Izergina N, Caussinus E, Reichert H. Amplification of neural stem cell proliferation by intermediate progenitor cells in Drosophila brain development. Neural Dev. 2008;3:5.

7. Turner-Evans DB, Jayaraman V. The insect central complex. Curr Biol. 2016 Jun 6;26(11):R453–7.

8. Syed MH, Mark B, Doe CQ. Steroid hormone induction of temporal gene expression in Drosophila brain neuroblasts generates neuronal and glial diversity. Elife [Internet]. 2017 Apr 10;6. Available from: https://www.ncbi.nlm.nih.gov/pubmed/28394252

9. Ren Q, Yan CP, Liu Z, Sugino K, Mok K, He Y, et al. Stem cell intrinsic, Seven-up-triggered temporal factor gradients diversify intermediate neural progenitors. Curr Biol. 2017;in press.

10. Bayraktar OA, Doe CQ. Combinatorial temporal patterning in progenitors expands neural diversity. Nature. 2013 Jun 19;498:445–55.

11. Abdusselamoglu MD, Eroglu E, Burkard TR, Knoblich JA. The transcription factor odd-paired regulates temporal identity in transit-amplifying neural progenitors via an incoherent feed-forward loop. VijayRaghavan K, Wang H, Sen S, editors. eLife. 2019 Jul 22;8:e46566.

12. Tang JLY, Hakes AE, Krautz R, Suzuki T, Contreras EG, Fox PM, et al. NanoDam identifies Homeobrain (ARX) and Scarecrow (NKX2.1) as conserved temporal factors in the Drosophila central brain and visual system. Dev Cell. 2022 May 9;57(9):1193–1207.e7.

13. Álvarez JA, Díaz-Benjumea FJ. Origin and specification of type II neuroblasts in the Drosophila embryo. Dev Camb Engl. 2018 Apr 5;145(7):dev158394.

14. Walsh KT, Doe CQ. Drosophila embryonic type II neuroblasts: origin, temporal patterning, and contribution to the adult central complex. Dev Camb Engl. 2017 Dec 15;144(24):4552–62.

15. Chell JM, Brand AH. Nutrition-responsive glia control exit of neural stem cells from quiescence. Cell. 2010 Dec 23;143(7):1161–73.

16. Lai SL, Doe CQ. Transient nuclear Prospero induces neural progenitor quiescence. Elife. 2014;3.

17. Maurange C, Cheng L, Gould AP. Temporal transcription factors and their targets schedule the end of neural proliferation in Drosophila. Cell. 2008 May 30;133(5):891–902.

18. Bayraktar OA, Boone JQ, Drummond ML, Doe CQ. Drosophila type II neuroblast lineages keep Prospero levels low to generate large clones that contribute to the adult brain central complex. Neural Dev. 2010;5:26.

19. Yu HH, Awasaki T, Schroeder MD, Long F, Yang JS, He Y, et al. Clonal development and organization of the adult Drosophila central brain. Curr Biol. 2013 Apr 22;23(8):633–43.

20. Yang JS, Awasaki T, Yu HH, He Y, Ding P, Kao JC, et al. Diverse neuronal lineages make stereotyped contributions to the Drosophila locomotor control center, the central complex. J Comp Neurol. 2013 Aug 15;521(12):2645–62.

21. Yang CP, Samuels TJ, Huang Y, Yang L, Ish-Horowicz D, Davis I, et al. Imp and Syp RNA-binding proteins govern decommissioning of Drosophila neural stem cells. Dev Camb Engl. 2017 Oct 1;144(19):3454–64.

22. Samuels TJ, Järvelin AI, Ish-Horowicz D, Davis I. Imp/IGF2BP levels modulate individual neural stem cell growth and division through myc mRNA stability. eLife. 2020 Jan 14;9:e51529.

23. Liu Z, Yang CP, Sugino K, Fu CC, Liu LY, Yao X, et al. Opposing intrinsic temporal gradients guide neural stem cell production of varied neuronal fates. Science. 2015 Oct 16;350(6258):317–20.

24. Toledano H, D’Alterio C, Czech B, Levine E, Jones DL. The let-7-Imp axis regulates ageing of the Drosophila testis stem-cell niche. Nature. 2012 May 23;485(7400):605–10.

25. Wu YC, Chen CH, Mercer A, Sokol NS. Let-7-complex microRNAs regulate the temporal identity of Drosophila mushroom body neurons via chinmo. Dev Cell. 2012 Jul 17;23(1):202–9.

26. Riebli N, Viktorin G, Reichert H. Early-born neurons in type II neuroblast lineages establish a larval primordium and integrate into adult circuitry during central complex development in Drosophila. Neural Dev. 2013;8:6.

27. Liu J, Liu W, Yang L, Wu Q, Zhang H, Fang A, et al. The Primate-Specific Gene TMEM14B Marks Outer Radial Glia Cells and Promotes Cortical Expansion and Folding. Cell Stem Cell. 2017 Nov 2;21(5):635–649.e8.

